# Assessment of antimicrobial activity of insects’ products and nests used in traditional medicine in Burkina Faso

**DOI:** 10.1101/2023.11.25.568644

**Authors:** Mamadou Ouango, Hama Cissé, Rahim Romba, Samuel Fogné Drabo, Rasmané Semdé, Aly Savadogo, Olivier Gnankiné

**Affiliations:** Laboratoire d’Entomologie Fondamentale et Appliquée, Université Josep KI ZERBO, 03 BP 7021, Ouagadougou, Burkina Faso; Laboratoire de Biochimie et Immunologie Appliquées, Université Joseph KI ZERBO, 03 BP 7021, Ouagadougou, Burkina Faso; Laboratoire du Développement du Médicament, Centre de Formation, de Recherche et d’Expertise en Sciences du Médicament, Université Joseph KI ZERBO, 03 BP 7021, Ouagadougou, Burkina Faso

**Keywords:** Insect products, Termite nests, Antimicrobial activity, Pathogen strain, Traditional medicine, Burkina Faso

## Abstract

In Burkina Faso, products and insect nests are used for therapeutic purposes in traditional medicine. However, this use by local populations is marginal and empirical. Our study aimed at evaluating the antimicrobial activity of insect products and their nests. For this purpose, the collected insect products and nests were finely ground. Hydroethanolic extraction of bioactive molecules with potential antibacterial activity was performed according to standard methods. The solid medium diffusion method was used to test the antibacterial activity of hydroethanolic extracts of honey bee, bee wax, propolis, and termite nests against 22 pathogenic strains by inhibition diameter. Imipenem was used as a positive control. The extraction yields varied from 7.33% to 35.38% depending on the content of soluble matter. All products extracts and insect nests tested showed inhibitory activities. The inhibition diameters varied depending on the extract and strain tested. The largest diameter of inhibition (26±0.0 mm) was obtained using the nest extract of *Macrotermes bellicosus* against *Salmonella* Typhimurium ATCC14028. The lowest diameter of inhibition was 07±0.0 mm obtained with honey extract against Pseudomonas aeruginosa ATCC27853. Index multi-resistance of the extracts tested were between 0.2 and 0.6. Interestingly, the inhibition diameters of certain products and nest extracts of insects were sometimes greater than those of imipenem against the strains tested. This study revealed the antimicrobial potential of termite nest extracts and hive products against pathogens.

## Introduction

Humans use insects and their products for various purposes [1]. Insects and their products are sometimes used in the diet of both humans [2–4] and farm animals [5, 6]. They are also used to feed on insects [7]. Insects and their products are also used in different forms and preparations for treating pathologies, particularly in traditional medicine around the world [8, 9]. Some traditional therapeutic practices persist and co-exist with those of modern medicine in treating various pathologies. Thus, maggot therapy remains used in most hospitals [10] for the treatment of ulcerating wounds. Similarly, honey is increasingly being used for the treatment of burns and wound healing [11]. The use of termite mounds in traditional societies for treating traumatic and diarrheal pathologies is still current [12–14]. Indeed, in 2022, a study conducted in Burkina Faso by Ouango et al. (2022) restricted to traditional health practitioners revealed the use of wax, honey, propolis, and honeycomb (*Macrotermes bellicosus* and *Trinervitermes* sp) for the production of traditional remedies offered to patients [15]. However, data on the antimicrobial effects of these products and insect nests on pathogenic germs remain scarce.

There is no doubt that antibiotics provide efficacy in many bacterial diseases if diagnostics are well established by medicinal practitioners. Until now, antibiotics have been used to treat infectious diseases. Unfortunately, their use has sometimes been accompanied by the emergence of resistant strains that threaten their effectiveness [16]. Antimicrobial resistance occurs when germs, such as bacteria and fungi, develop the ability to defeat the drugs designed to kill them. Resistant infections can be difficult and sometimes impossible to treat. With regard to the increasing antibiotic resistance to bacteria associated with auto-medication practiced by many people in Africa, many researchers have focused on the search for alternative antibiotic-resistant strains.

Currently a global concern, antimicrobial resistance (AMR) poses a threat to human, animal, and environmental health. However, the Food and Agriculture Organization of the United Nations recently described AMR as a quintessential one-health problem. The major driver of antimicrobial resistance is the inappropriate use of antimicrobials in human health care and animal production [17]. However, the use of fractions with potential antibacterial activity (FPAA) from products and insect nests could be an alternative to skirt antibioresistance. Therefore, our study aimed to evaluate the antimicrobial activity of wax, honey, propolis, and nests of *Macrotermes bellicosus* and *Trinervitermes* sp.

## Materials and Methods

### Insects’ collection sites

Product extracts and insect nests were collected in three provinces (Fig. 1) belonging to three different phytogeographical zones, namely Kadiogo (12°21′ 56.4′′N; 1°32′2′′ W), Houët (11 °7′ 55.36′′N; 4°14′0.01′′ W), and Séno (14°1′ 48 ′′N; 0°1′48′′ W). These sites were where we conducted a survey of traditional healers to collect information on the use of medicinal insects. These insect collection sites were chosen for their cosmopolitan character, being the largest provinces of identified climatic zones, and their great potential as traditional healers.

**Fig. 1:**
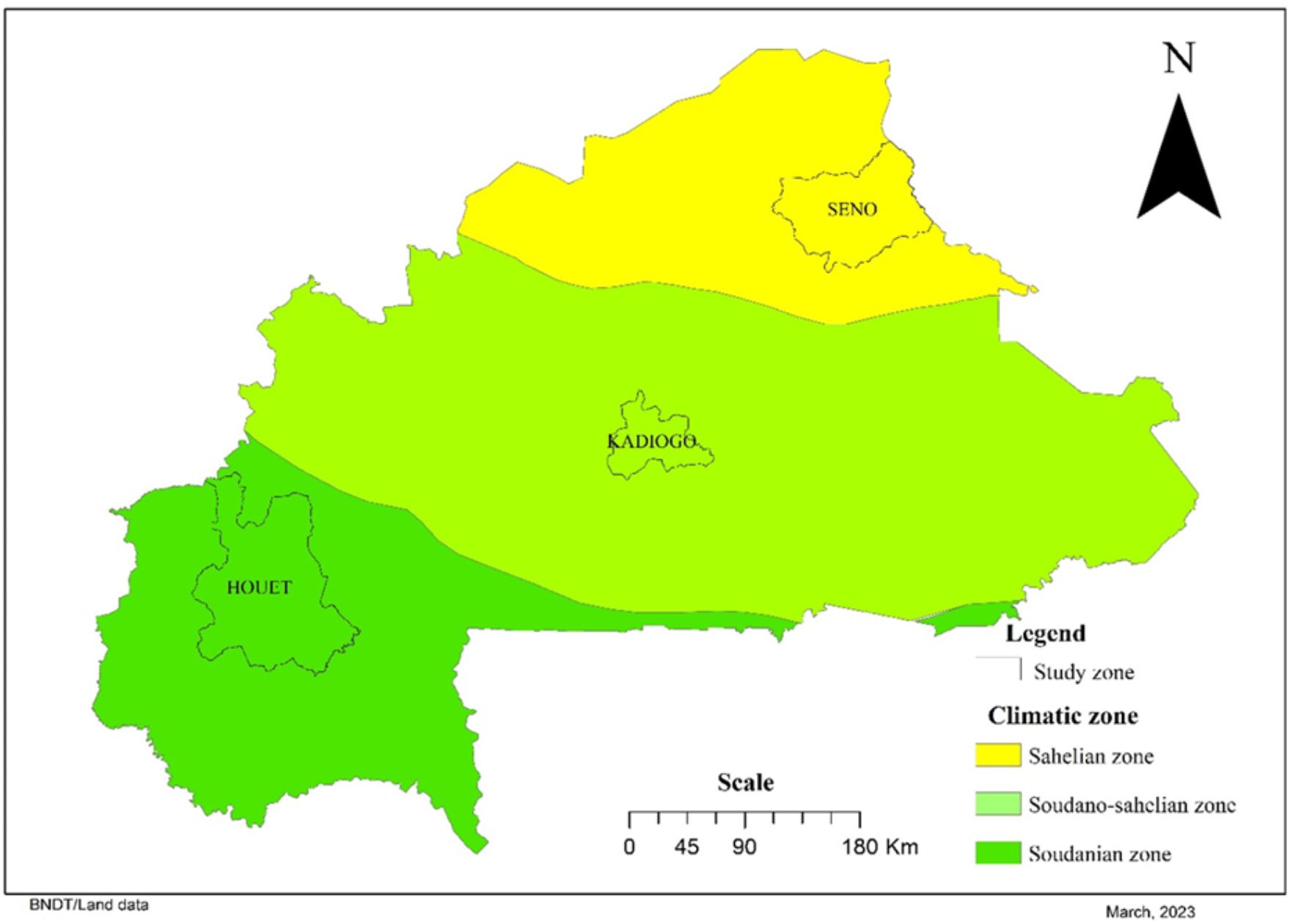
Location of insect sample collection sites.

### Sampling of insect products and nests insect

Three insect products and two insect nests were collected for this study. These products were collected on the basis of information provided according to traditional healers (Fig. 2). These are wax, honey, and bee propolis from *Apis mellifera* collected from beekeepers and termite nests of *Macrotermes bellicosus* and *Trinervitermes* sp. collected from the wild during outings at the study sites. The propolis was collected at Bobo Dioulasso (Houët province). Honey and wax were collected at Gonsé (Kadiogo province). Nest fragments were collected at Dori (Séno province). These products were packaged in sterile jars and transported to the laboratory for future use.

**Fig. 2.**
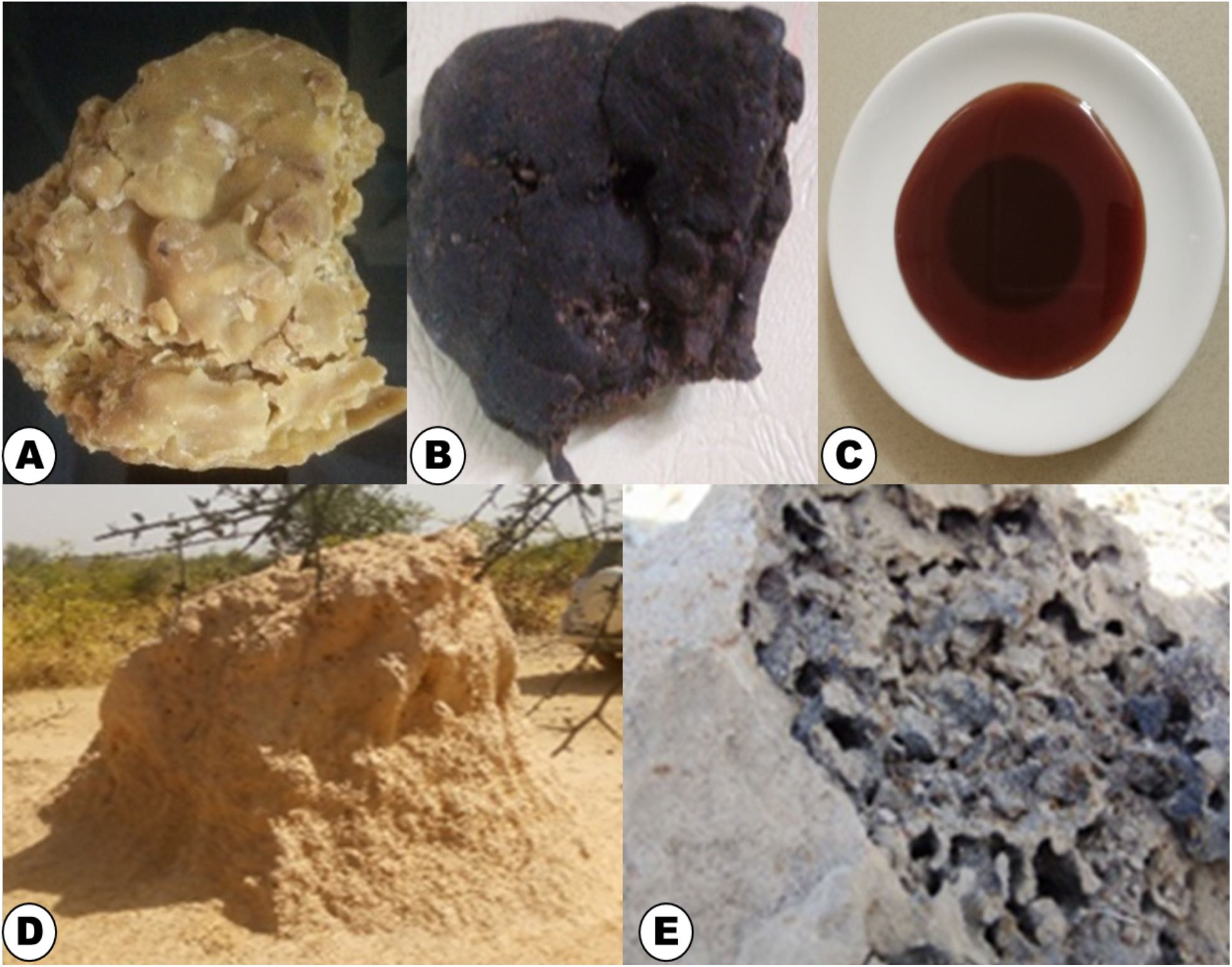
Bee products and insect nests collected. A : Bee’s wax (Gonse); B : Bee propolis (Bobo Dioulasso); C : Honey (Gonse); D : Nest of *Macrotermes bellicosus* (Dori); E : Nest of *Trinervitermes* sp (Dori).

### Extraction of FPAA from products and nests collected

Fragments of each nest were finely ground using a laboratory mortar. Bee wax and bee propolis were finely fragmented. The powders and fine fragments obtained were packaged in Falcon tubes (CONICAL BOTTOM CELLSTAR® STERILE). The extraction of FPAA was performed according to the method described by Dah-Nouvlessounon *et al.* [18], which has been readapted. Thus, for insect nests (aqueous extracts), aqueous extraction of FPAA was performed using sterile ultrapure Milli-Q water. As for insect products (hydroethanolic extracts), hydroethanolic extraction to obtain the FPAA was performed from a sterile hydroalcoholic solution at 30% (ethanol and sterile Milli-Q ultrapure water in the proportions 30:70 v/v). Thus, 1g of each shredded product and insect nest was macerated in 10 mL of the extraction solution for 12 h at 25 °C under magnetic stirring with sterile Milli-Q ultrapure water and sterile hydroalcoholic solution, respectively. The macerates were centrifuged at 3,000 rpm for 10 min at 4 °C using a JOUAN BR4 refrigerated centrifuge. After centrifugation, the supernatants were collected in Eppendorf tubes and kept cool at 4 °C. After collecting the supernatants, the extraction solvents were evaporated to dryness in an oven at 45 °C until a dry extract of constant mass was obtained for the evaluation of extraction yield. The residues obtained were kept at 4 °C until the antimicrobial tests were performed.

### Extraction yield

The extraction yield was determined by the ratio between the mass of the powdered product or insect nest after extraction and the mass of the ground material at the start according to the following formula:

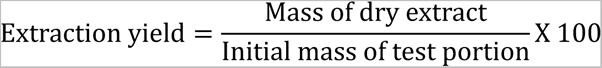

### Assessment of antimicrobial activity of FPAA from products and nests collected

#### Microbial strains used for antimicrobial testing

Antimicrobial activity tests were performed against 22 microbial germs, including 12 Gram negative bacteria (GNB), 7 Gram positive bacteria (GPB) and 3 fungal strains (Table 1). The microbial strains used in this study are based on several criteria: these strains are commonly of hospital and food origin because of their high incriminations in pathologies in animals and humans, and these strains are chosen based on their natural resistance to various types of antimicrobial agents.

**Table 1:** Microbial strains tested and their references.

| Microorganism | Species | Reference |
| --- | --- | --- |
| Gram-negative bacteria | *Escherichia coli* | 652654 |
|  | *Escherichia coli* | ATCC25922 |
|  | *Escherichia coli* | ATCC8739 |
|  | *Klebsiella pneumoniae* | 203 |
|  | *Klebsiella pneumoniae* | ATCC13883 |
|  | *Providencia rettgeri* | 652655 |
|  | *Pseudomonas aeruginosa* | ATCC9027 |
|  | *Pseudomonas aeruginosa* | ATCC27853 |
|  | *Salmonella abony* | NCTC6017 |
|  | *Salmonella enteritidis* | ATCC13076 |
|  | *Salmonella Typhimurium* | ATCC14028 |
|  | *Serratia odorifera* | 652411 |
| Gram positive bacteria | *Staphylococcus aureus* | ATCC43800 |
|  | *Staphylococcus saprophyticus* | 652615 |
|  | *Clostridium perfringens* | ATCC13124 |
|  | *Clostridium sporogenes* | MAR0319404 |
|  | *Enterococcus faecalis* | ATCC29212 |
|  | *Bacillus pumilus* | ATCC14884 |
|  | *Listeria monocytogenes* | ATCC19112 |
| Fungal strains | *Saccharomyces cerevisiae* | ATCC9763 |
|  | *Yarrowia lipolytica* | W29 |
|  | *Candida shehatae* | ATCC22984 |

### Antimicrobial activity testing of FPAA from products and nests collected

The antimicrobial activity of FPAAs was tested according to the agar diffusion method described by Kirby-Bauer, following the guidelines of the Clinical Laboratory Standards Institute. The preparation of microbial inoculate was performed from young colonies aged 16– 18 h diluted in test tubes containing physiological saline. All microbial suspensions obtained were adjusted to a turbidity of 0.5 MacFarlant. The value of this standard turbidity of 0.5 McFarland corresponds approximately to a culture density of 1.5×10^8^ cells/mL. For the preparation of disks containing the FPAAs, blank and sterile test antibiogram discs (MASTDISCS® AST) of 6 mm of diameter were used. These discs were impregnated with solutions of extracts of insect products or insect nests contained in Eppendorf tubes for 10 min. For performing the antibiogram, Mueller– Hinton and Sabouraud chloramphenicol agar were used for the bacterial and fungal strains, respectively. Petri dishes containing the agars were inoculated by swab test with different bacterial and fungal strains. The inoculated agars were dried near a Bunsen burner for 5 min before receiving discs impregnated with product extracts and insect nests. Antibiotic discs (Imipenem) were used as positive controls, and blank and sterile test antibiogram disks impregnated in DMSO without extract were used as negative controls. After depositing the discs, the Petri discs were left at room temperature for 15 min to allow the diffusion of extracts and incubated at 37 °C for 24 h. The inhibition diameters materialized by a clear halo around the discs were measured using a BioNumerical ruler (MICROBIAL DATA ANALYSIS SOFT WARE).

### Determination of index multi-resistance to extracts of products and insect nests

Index of multi-resistance to extracts (IMRE) of products and nests of insects was determined according to Das *et al.* [19]. The microbial strains tested were classified according to IMRE. IMRE was calculated using the following formula:

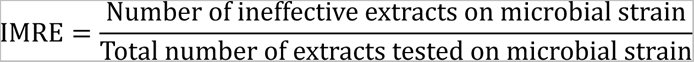

### Data processing and statistical analyses

Results were expressed as mean number followed by standard deviation (M± SD) and subject to Student’s t-test using R. The significance threshold was 5%. XLSTAT-2019 software was used for principal component analysis (PCA). PCA was used to explore the correlation between the activities of product and nest extracts of insect and the different pathogen strains.

## Results

### Yield values of extractions of the products and insect nests

The yields of the different extractions varied according to the products and nest insects used (Fig. 3). The best extraction yield (35.38%) was obtained using honey. The lowest yield (7.33%) was obtained from the nest of *Macrotermes bellicosus*.

**Fig. 3.**
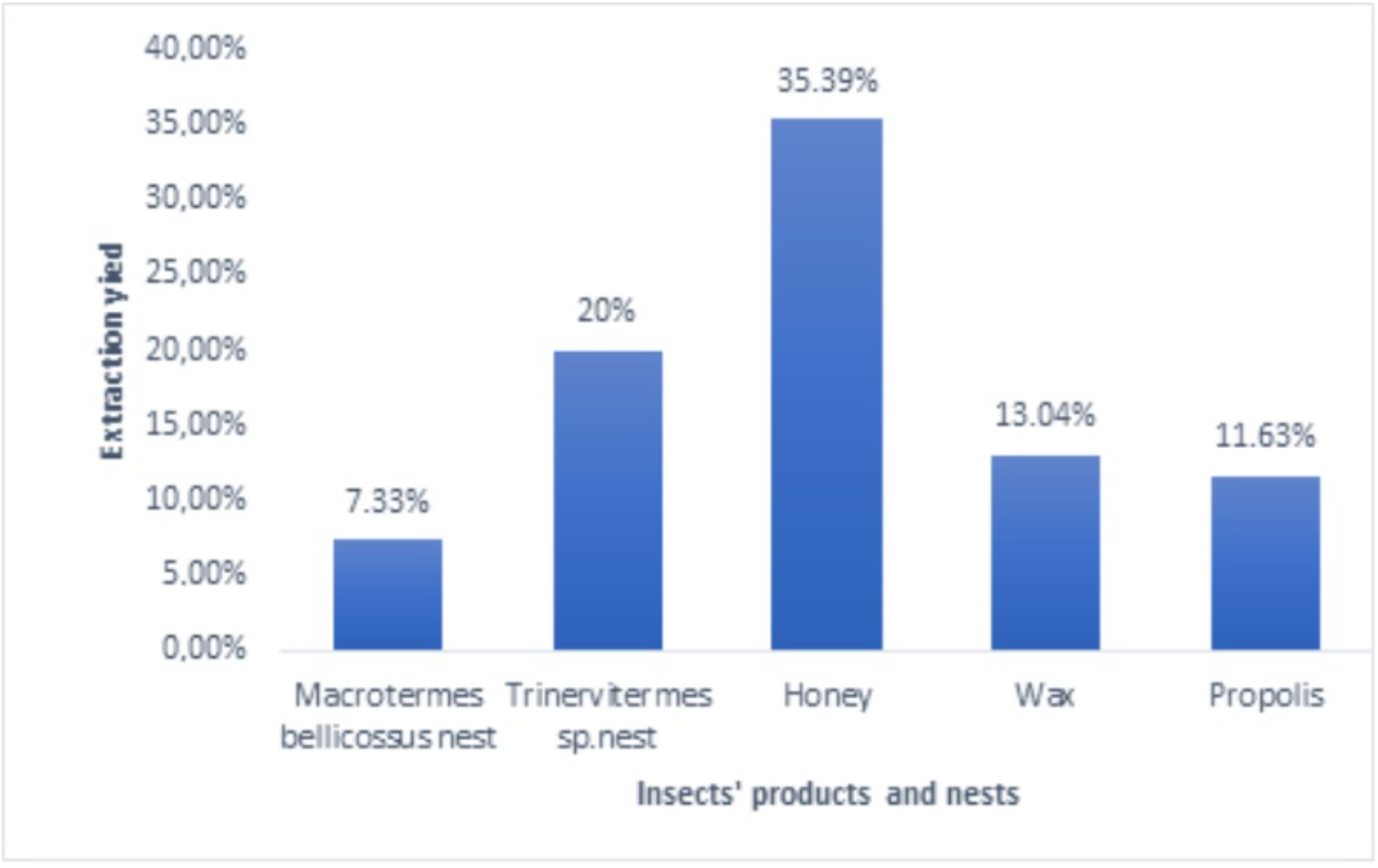
Extraction yield of the products and the nests of insects.

### Antimicrobial activity of FPAA of the aqueous and hydroethanolic extracts

The inhibition diameters of various FPAAs of the products and nest insects tested (Fig. 4) are shown in Table 2. These results indicated that the microbial strains were sensitive to the different extracts used. Inhibition diameters ranged from 07±00 mm to 26±00 mm against strains tested for FPAAs showing antibacterial activity. The DMSO-impregnated discs had no antimicrobial activity against any of the strains tested.

**Fig. 4.**
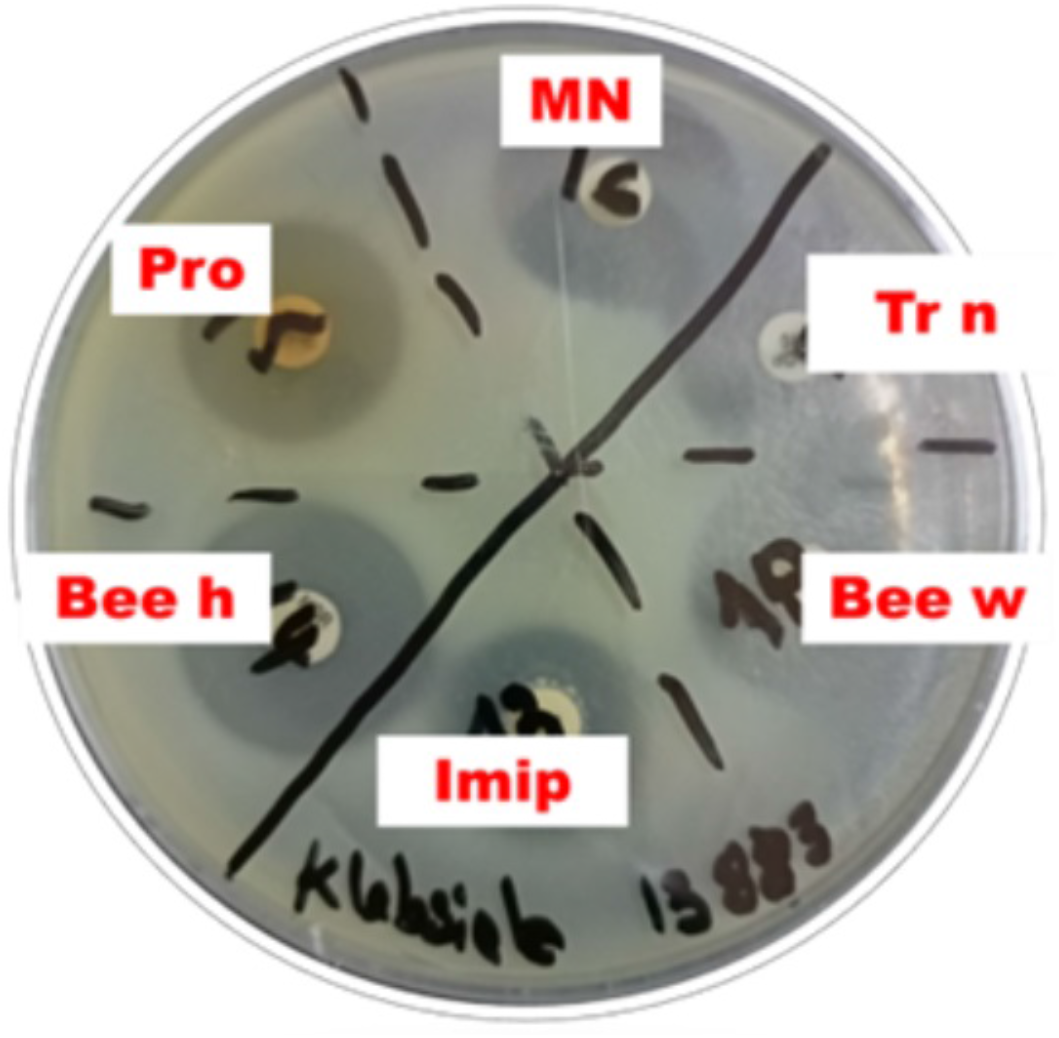
Antimicrobial activity of FPAA of the aqueous and hydroethanolic extracts on *Klebsiella pneumoniae* ATCC13883. Legend: Imip : imipenem; Bee h : Bee honey; Pro : Propolis; MN : Nest of *Macrotermes bellicosus*; Tr n : Nest of *Trinervitermes* sp.; Bee w: Bee wax

**Table 2.**
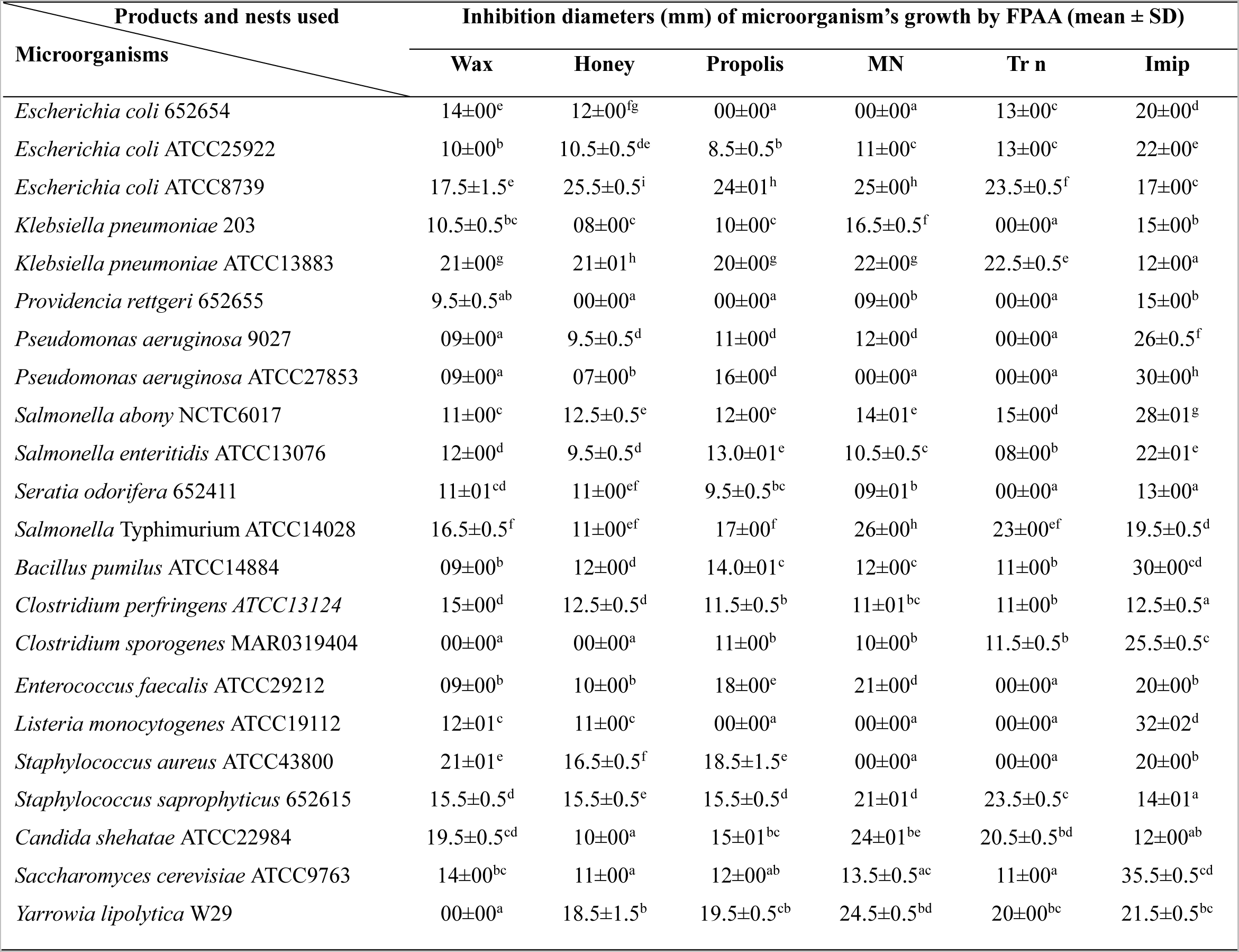
Antimicrobial activities of FPAA of the aqueous and hydroethanolic extracts used. SD: Standard Deviation; MN: Nest of *Macrotermes bellicosus*; Tr n: Nest of *Trinervitermes* sp.; Imip: imipenem. Values in same column with different superscript letters are significantly different (p < 0.05) according to Student’s t-test.

The hydroethanolic extract from wax was active against 20 of the 22 strains tested with an inhibition percentage of 91%. The largest inhibition diameter was obtained against *Klebsiella pneumoniae* ATCC13883 (21±00 mm) while that of smallest diameter of inhibition (09±00 mm) was found against two strains of *Pseudomonas aeruginosa*, *Bacillus pumilus* ATCC14884, and *Enterococucus faecalis* ATCC29212. However, the hydroalcoholic extract of wax had no inhibitory effect on *Clostridium sporogenes* MAR0319404 and *Yarrowia lipolytica* W29.

The hydroethanolic extract of honey was active in 91% of the microbial strains tested. However, no inhibition was observed on *Providencia rettgeri* 652655 and *Clostridium sporogenes* MAR0319404. The largest inhibition diameter was obtained against *E. coli* ATCC8739 (25±0.5 mm) and the smallest inhibition diameter (07±00 mm) on *Pseudomonas aeruginosa* ATCC27853.

The hydroalcoholic extract of propolis was active on 86.36% (19/22) of the bacteria tested. However, this extract had no antimicrobial activity against *E. coli* 652654, *Providencia rettgeri* 652655 and *Listeria monocytogenes* ATCC19112. The largest inhibition diameter (24±01 mm) was obtained against *E. coli* ATCC8739 and the smallest inhibition diameter (08±0.5 mm) was obtained against *E. coli* ATCC25922.

The aqueous extract from the nest of *Trinervitermes* sp. was active against of 14 strains out of the 22 strains tested (63.64%). *Pseudomonas aeruginosa* ATCC9027, *Pseudomonas aeruginosa* ATCC27853, *Klebsiella pneumoniae* 203, *Providencia rettgeri* 652655, *Seratia odorifera* 652411, *Enterococcus faecalis* ATCC29212, *Listeria monocytogenes* ATCC19112, and *Staphylococcus aureus* ATCC43800 strains tested were insensitive to aqueous extracts from the nest of *Trinervitermes* sp. However, the largest diameter (23.50±0.50 mm) was obtained against *Staphylococcus saprophyticus* 652615 and *E. coli* 652654.

The hydroethanolic extract of the nest of *Macrotermes bellicosus* was active on 18 of 22 microbial strains tested, i.e., an inhibition rate of 81.82%. The highest diameter of inhibition was 26±00 mm on *Salmonella Typhimurium* ATCC14028 compared with the diameter of inhibition reported with imipenem (19±00 mm). The lowest reported inhibition diameter was 09±00 mm against *Providencia rettgeri* 652655, whereas the extract was ineffective against *E. coli* 652654, *Pseudomonas aeruginosa* 27853, *Listeria monocytogenes* ATCC19112, and *Staphylococcus aureus* ATCC43800.

### Effectiveness of some extracts compared to imipenem

The inhibition diameters of extracts of the products and nest insects were sometimes greater than those of imipenem. Indeed, this observation was reported on 43.49% (10/23) of the microbial strains tested. Table 3 lists the microbial strains on which the extracts of the products and nest insects have greater inhibitory activity than that of imipenem. For these extracts, the smallest difference in inhibition diameters was 01 mm obtained with the nest extracts of *Macrotermes bellicosus* and the extract of wax on *Enterococucus faecalis* ATCC29212 and *Staphylococcus aureus* ATCC43800. The largest difference of inhibition diameter was 12 mm obtained using an extract of the nest of *Macrotermes bellicosus* versus *Candida shehatae* ATCC22984. Note that the extract from the nest of *Macrotermes bellicosus* inhibited the growth of the greatest number of these strains (70% of strains).

**Table 3.** Extracts of products and nests insects showing inhibition diameter greater than imipenem.

| Microorganisms | Products and nests insects |
| --- | --- |
| *Escherichia coli* ATCC8739 | Wax, honey, propolis, nest of *Macrotermes bellicosus* and nest of *Trinervitermes* sp. |
| *Klebsiella pneumoniae* 203 | Nest of *Macrotermes bellicosus* |
| *Klebsiella pneumoniae* ATCC13883 | Wax, honey, propolis, nest of *Macrotermes bellicosus* and nest of *Trinervitermes* sp. |
| *Salmonella* Typhimurium ATCC14028 | Nest of *Macrotermes bellicosus* |
| *Clostridium perfringens* ATCC13124 | Wax |
| *Enterococcus faecalis* ATCC29212 | Nest of *Macrotermes bellicosus* |
| *Staphylococcus aureus* ATCC43800 | Wax |
| *Staphylococcus saprophyticus* 652615 | Wax, honey, propolis, nest of *Macrotermes bellicosus* and nest of *Trinervitermes* sp. |
| *Candida shehatae* ATCC22984 | Wax, propolis and nest of *Macrotermes bellicosus* |
| *Yarrowia lipolytica* W29 | Nest of *Macrotermes bellicosus* |

### IMRE of products and nests of insects

The IMREs of the microbial strains used to extract the products and nest insects are shown in Fig. 5. These indices ranged from 0% to 60%. Analysis of these IMRE of products and nests of insects revealed that some microbial strains were very sensitive to the extracts tested (0%). However, there were strains that were more or less sensitive to the extracts tested (20% to 60%). Concerning Gram-negative bacteria, 6 strains (*Escherichia coli* ATCC25922, *Escherichia coli* ATCC8739, *Klebsiella pneumoniae* ATCC13883, *Salmonella abony* NCTC6017, *Salmonella enteritidis* ATCC 13076 and *Salmonella typhimurium* ATCC14028) were sensitive to all extracts tested. For Gram-positive bacteria, three strains, *Bacillus pumilus* ATCC14884, *Clostridium perfringens* ATCC13124, and *Staphylococcus saprophyticus* 652615, were sensitive to all extracts of products and nest insects. On the other hand, for the fungal strains, two strains, Candida shehatae ATCC22984 and *Saccharomyces cerevisiae* ATCC9763, were sensitive to all extracts tested. Thus, the most resistant strains to these extracts of products and nest insects with an IMRE value of 0.60 were obtained with *Providencia rettgeri* 652655, *Listeria monocytogenes* ATCC19112 and *Staphylococcus aureus* ATCC43800.

**Fig. 5.**
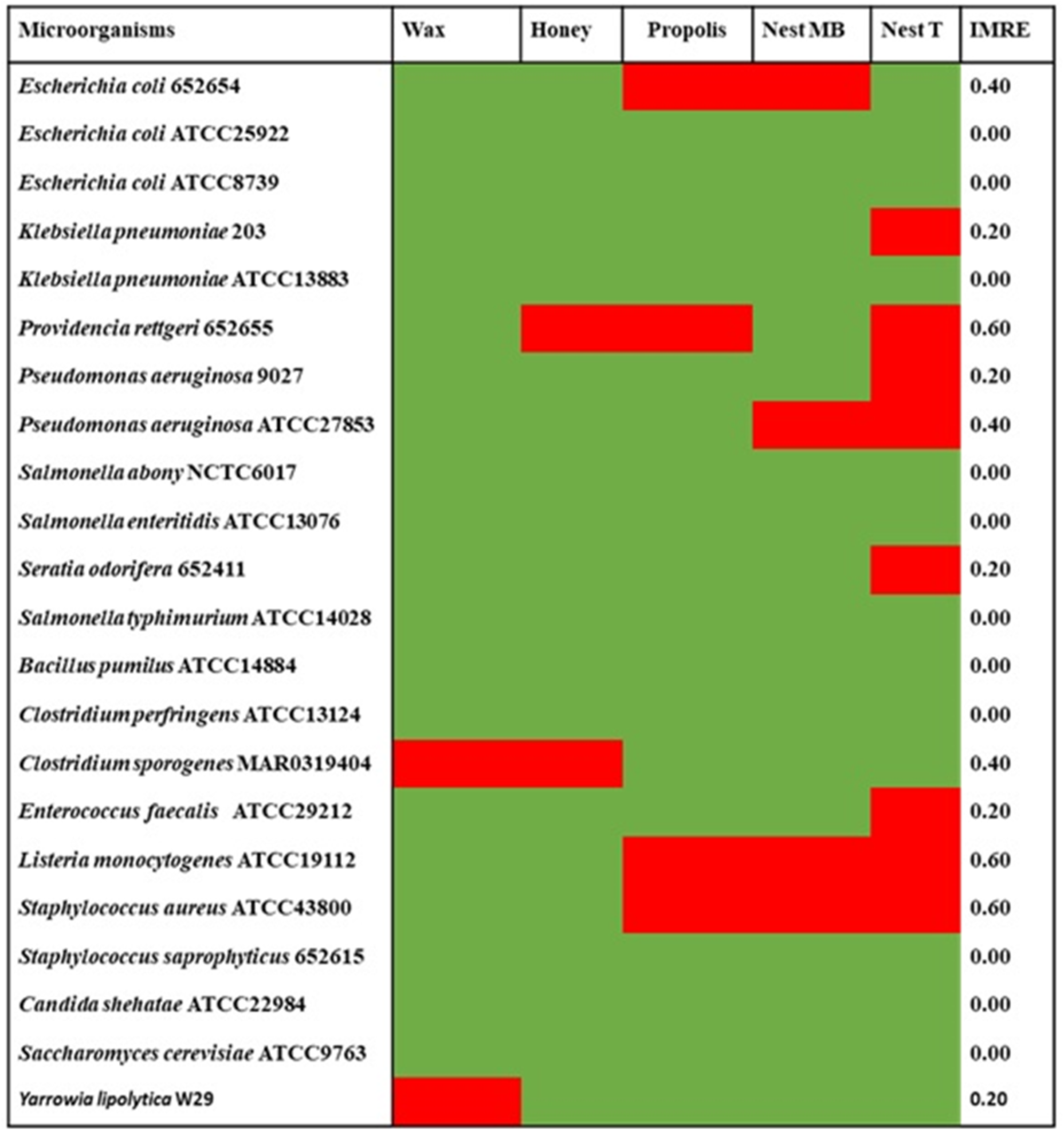
IMRE values of bacterial strains to extracts of products and nests of insects; Nest MB: Nest of *Macrotermes bellicosus*; Nest T: Nest of *Trinervitermes* sp.; Red: Resistant; Green: sensitive

Strains less resistant to extracts of products and nest insects with a low IMRE of 0.40 were *Escherichia coli* 652654, *Pseudomonas aeruginosa* ATCC 27853, and *Clostridium sporogenes*. The IMRE value of 0.20 for extracts of products and nest insects was found with strains *Klebsiella pneumoniae* 203, *Pseudomonas aeruginosa A*TCC9027, *Seratia odorifera* 652411, *Enterococcus faecalis* ATCC29212, and *Yarrowia lipolytica* W29. Overall, 11 microbial strains tested had a value 0.00 for IMRE.

### Principal component analysis between FPAA and microbial strains

The first two axes (F1 and F2) explain 76.73% of the dispersion of variables, which are bacterial and fungal strains (Fig. 6.). PCA shows a strong positive correlation between termite nest extracts, propolis, and some microbial strains. These are *Yarrowia lipolytica, Salmonella* Typhimurium, *and Staphylococcus saprophyticus*. Hydroalcoholic extracts from termite nests and propolis therefore have a broad spectrum of action. However, the extract from the nest of *Macrotermes bellicosus* is much more effective in inhibiting the growth of *Yarrowia lipolytica* W29 than other extracts, given the strong correlation between these two parameters. The wax and honey of Apis mellifera exhibit high inhibitory actions are high against *Staphylococcus aureus* ATCC43800, *Klebsiella pneumoniae* ATCC13883 and *Escherichia coli* ATCC8739. However, the best growth inhibition of *Staphylococcus aureus* ATCC43800 was achieved with the wax extract.

**Fig. 6.**
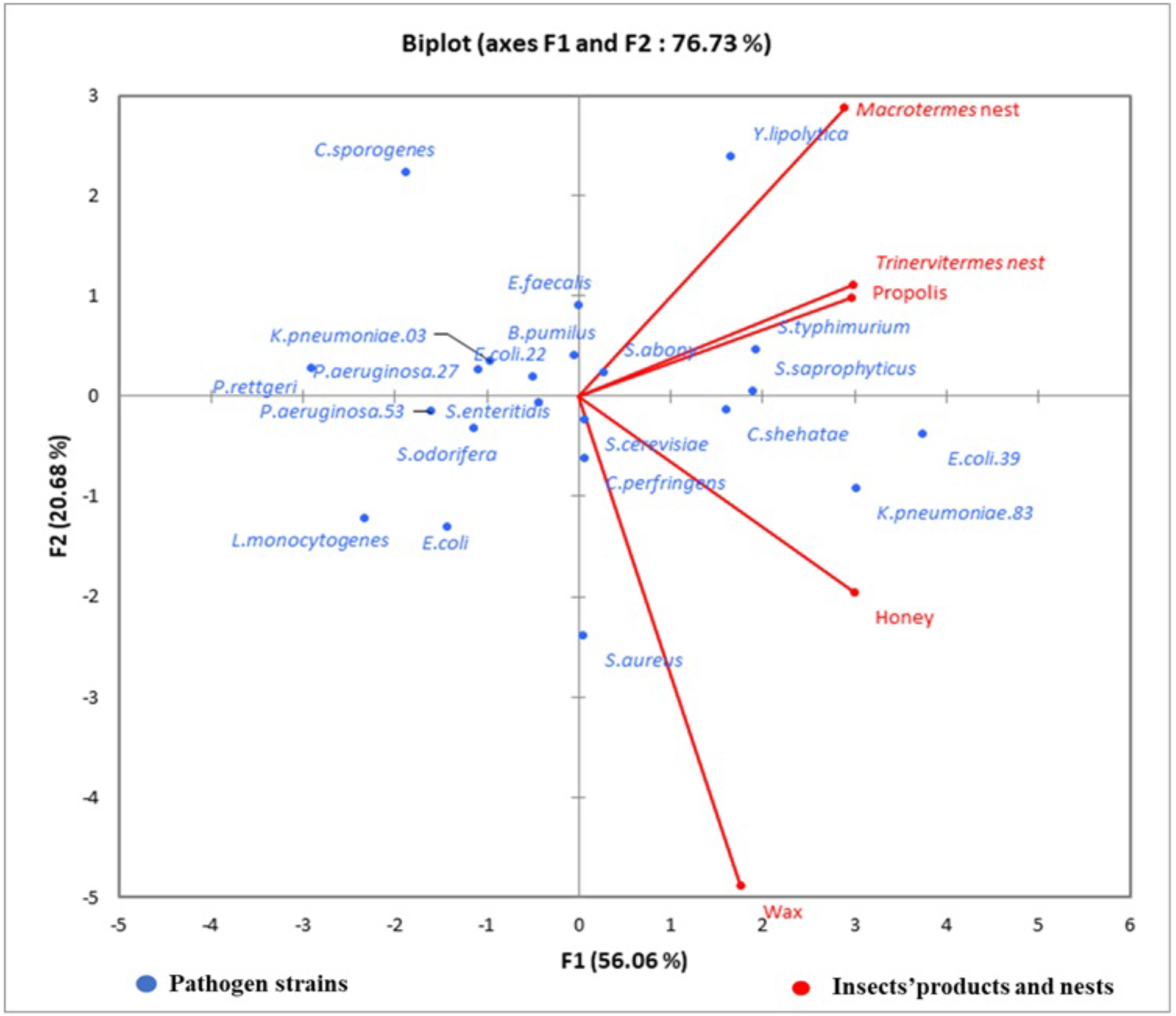
Distribution of the products and nests of insects to microbial strains according to the PCA plan

## Discussion

Increasing antimicrobial resistance is a serious global issue that has steered research for the identification of new biomolecules with broad antibacterial activity. The insects and their derivatives, such as products and nests, are often used in traditional medicine. In nature, the biomolecules produced by insects play an important role in their protection. The products and nests of insects contain a wide variety of biomolecules capable of inhibiting or slowing the growth of bacteria. Thus, honey contains a more soluble matter in the hydroalcoholic solution, unlike the nest of *Macrotermes bellicosus*, which has a lower yield. The biomolecules of products and nests of insects may have variable mechanisms of action affecting the cell membrane and cytoplasm, and in some cases can completely change cell morphology, even the expression of genes (Fig. 7).

**Fig. 7.**
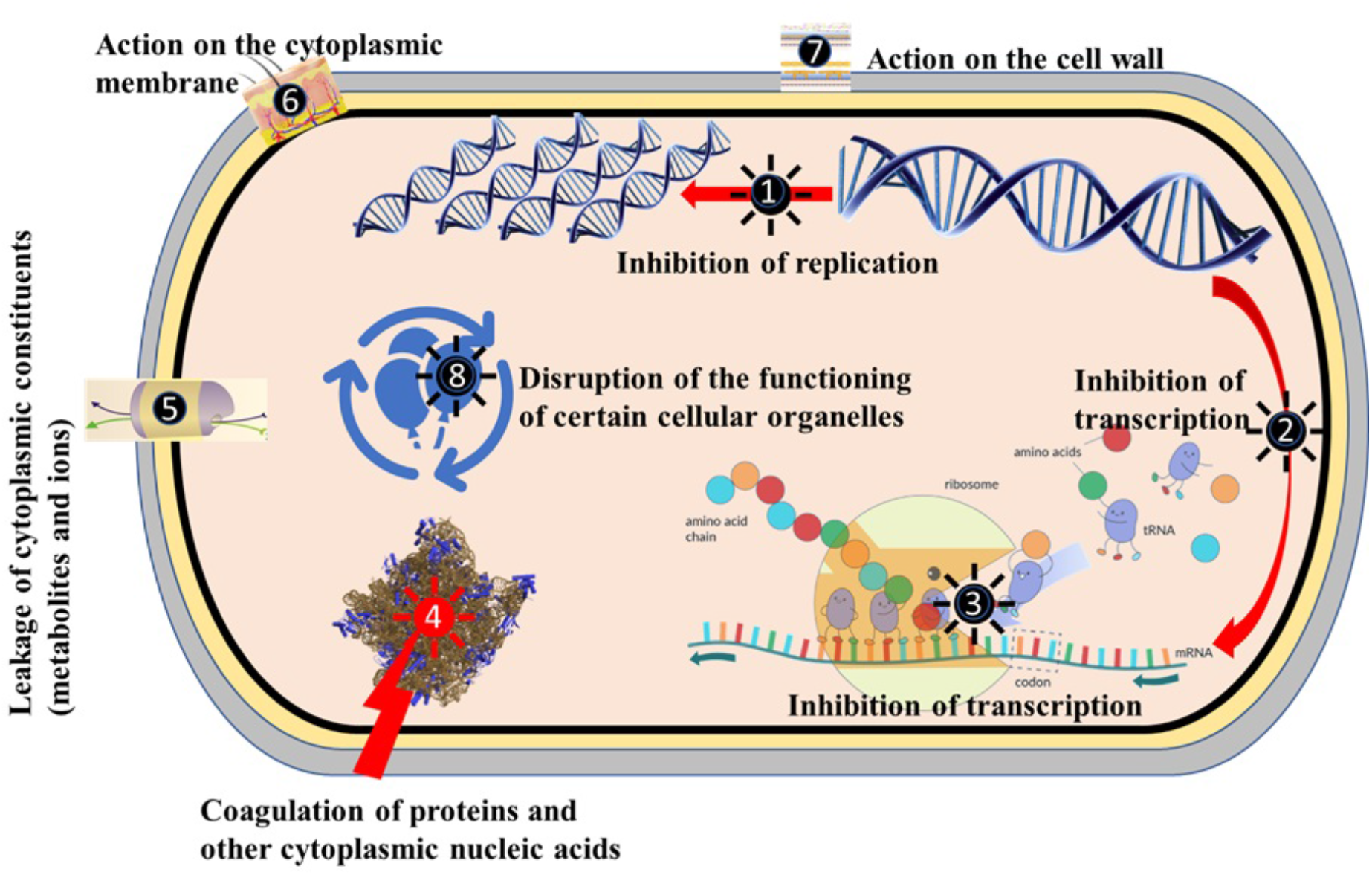
Putative mechanisms of action of the extracts of products and nests insects on microbial cells

The antimicrobial activity of wax has long been empirically exploited by several people for treating certain pathologies in humans. Costa-Neto [20] reported the use of wax by Siribinha fishermen in America to treat ear infections. Wax is a mixture of approximately 300 molecules, including monoesters, diesters, triesters, hydrocarbons, hydroxy mono- and polyesters, free alcohols, and other unidentified substances [21]. Some of these molecules have shown inhibitory activities against different pathogenic microbial [22]. Thus, Octanal contained in wax and isolated from other natural products showed antimicrobial activity against *E. coli*, *Staphylococcus aureus* [23, 24], and *Listeria monocytogenes* [24, 25]. The results reveal antibacterial activity against the strains mentioned above. The antibacterial activity of wax and propolis against *Salmonella* is due to the presence of limonene in these products [26, 27]. As for the anti-salmonella activity revealed, it would be due to the wax dodecanal [28]. Kinderlerer and Lund [29] demonstrated the antibacterial activity of hexanoic acid contained in wax against *Listeria monocytogenes*. Inouye et al. [23] also reported the inhibitory effect of geraniol, another wax component, against *E. coli* and *Staphylococcus aureus*.

Honey contains more than 150 compounds, including fructose (38%), sucrose (17%) and glucose (31%) and trisaccharides, 4% acid (gluconic acid), minerals, oxygenated water, proteins, enzymes, amino acids, vitamins, aromatic organic molecules, lipids and microscopic elements such as pollens, spores, algae, and yeasts [1]. The antimicrobial activity of honey is due to four factors: low water content (15-21%), oxygenated water content, acidity (pH=3.9) linked to gluconic acid, and the presence of organic molecules with antibiotic properties linked to the floral origin of honey [30].

The bactericidal activity of propolis is fairly well known with a broad spectrum of action on strains of *Staphylococcus* (*aureus* and *mutans*), *Streptococcus* (*mutans* and *sanguinis*), *Bacillus* (*cereus* and *subtilis*), *Proteus* (*vulgaris* and *mirabilis*), *Pseudomonas*, *Listeria*, *Salmonella*, *Clostridium*, *Escherichia coli*, *Enterococcus*, and *Helicobacter pylori* [31]. The antifungal activity of propolis against many yeasts has also been revealed by various tests on several species of *Candida*, *Saccharomyces*, and *Cryptococcus*, as well as on species of *Aspergillus*, *Microsporum*, and *Trichophyton* (dermatophytes) [32]. Indeed, the hydroalcoholic extract of propolis was active on all fungal strains tested, confirming its broad spectrum of activity on fungal strains, which corroborates the results reported in this study. Propolis contains 50%–55% resins and balms, 30% waxes and fatty acids, 10% essential oils, 5% pollen, and 5% organic and mineral substances [33, 34]. However, its compounds are essentially phenolic substances with interesting biological activities [35]. These phenolic compounds are essentially flavonoids and phenolic acids. These compounds are endowed with antimicrobial activity. The most effective flavonoids are galangin, pinocembrin, pinostrobin, quercetin, and rhamnetin, and caffeic and ferulic acids are found among these acids [1]. The antimicrobial molecules of propolis can act directly against bacteria and fungi or suppress microbial virulence factors, such as anti-biofilm factors [36]. The biological activities of propolis, such as antimicrobial, antioxidant, immunomodulatory, analgesic, and tissue-regenerating activities, make this product the most commonly used among *Apis Mellifera* products [37]. In modern medicine, propolis is used to treat respiratory infections, eczema, wound healing, cancer, and to strengthen the immune system [38]. Propolis could be used as an alternative to microbial strains that are multi-resistant to antibiotics [39].

The antimicrobial activity of *Trinervitermes* sp. extracts could be due to the presence of trinervitanes, which are known to inhibit the development of microorganisms [1]. Secretions of *Trinerviterme*s soldiers contain lethal toxicants [40]. Certain genera of termites, particularly *Trinervitermes,* secrete several chemical groups, including terpenes (sesquiterpenes, monoterpenes, and diterpenes) and cyclic lactones [1, 41]. Trivervitans and deterpenes of these termites have proven antimicrobial and antifungal activities [41–44]. The Australian company Entocosm has even filed a patent for the antibiotic activity of Australian termite trinervitanes [1].

The antimicrobial activity of extracts of *Macrotermes* and their nests was also reported by Solavan et al. (2007) against *Escherichia coli*, *Pseudomonas putida*, and *Klebsiella* sp. [45]. Insect nests are exposed to both bacterial and fungal attacks; therefore, they induce nests with the antimicrobial compounds that they develop [46]. Termites produce antimicrobial peptides known as termicin, which have both antibacterial and antifungal properties [47]. Hammoud Mahdi et al. [48] also reported the production of hydroquinone, an antibacterial substance by *Macrotermes*. Sawadogo et al. [49] reported that several typical bacteria and actinobacteria isolated from *Macrotermes bellicosus* termite mound materials are good producers of antibacterial and antifungal substances.

## Conclusion

This study revealed the antimicrobial activity of extracts from insect nests and hive products. These extracts acted equally well against Gram-positive and Gram-negative bacteria and fungal strains. The results of this study can lead to further investigations to isolate and identify new natural bioactive molecules from nests and insect products to provide a therapeutic alternative to the phenomenon of bacterial multiresistance. It provides scientific knowledge on the use of certain insect nests and products by traditional health practitioners for treating their patients.

## Author contribution statement

All authors listed have significantly contributed to the development and the writing of this article.

## Data availability statement

Data included in article/referenced in article.

## Additional information

No additional information is available for this paper.

